# Monitoring Lake Okeechobee Harmful Algal Bloom populations and dynamics with a long-duration Autonomous Surface Vehicle

**DOI:** 10.1101/2023.08.23.554529

**Authors:** Veronica Ruiz Xomchuk, Scott Duncan, Malcolm N. McFarland, Jordon S. Beckler

## Abstract

This article describes the main findings of a full year of continuous operation of a 2-meter Autonomous Sail and Solar Surface Drone, the Nav2 (Navocean Inc.), as part of a Harmful Algal Bloom (HAB) monitoring program in Lake Okeechobee. The Nav2 was equipped with a set of water quality and atmospheric sensors, that recorded high frequency measurements (¡ 1 min) and transmitted near real-time information to allow reporting through a web portal for assessment and operation responses. Major findings include detection of HABs early in the year through chlorophyll (chl-a) and phycocyanin (phyco) fluorometric measurements, as well as different spatial scales of variability in the algal patches. The 24/7 high resolution monitoring allowed detection of patch motion and discrimination between growth and motion along a transect. Furthermore, the platform can potentially fingerprint specific HAB species based on the relatively fine-scale spatial expression of the phyco to chl-a ratio, which essentially captures the bloom macrostructure (e.g. surface scums versus more uniform sub-surface waves over 0.1 - 1 km scale). Sensor outputs, when converted to concentrations based on calibrated with pure laboratory standards, did not accurately yield true chl-a or phyco values when compared to validation samples, likely due to the high turbidity of the lake. However, routine solid-state validations of fluorometric measurements proved useful for assessing consistency in optical sensors to check for sensor drift (e.g. to due biofouling), which was not significant. Overall this demonstration shows that the Nav2 can uniquely and reliably provide in situ HAB and environmental monitoring capabilities in a large, turbid, shallow lake. We envision that platform as an innovative technology for water resource managers by providing turn-key long-duration baseline environmental data (hands-off waypoint navigation), early warnings of HABs for protecting human health, and for HAB mitigation monitoring.

## 1 Introduction

The global proliferation of Harmful Algal Blooms (HABs) in aquatic environments is encouraging the implementation of effective monitoring systems that can be used to better understand population dynamics and allow rapid response for mitigating ecosystem, human, and economic health impacts (Babin et al, 2008). Traditional methods for monitoring HAB dynamics are constrained in their ability to capture biological and environmental data at both relevant spatial and temporal scales (Cullen, 2008; Ruddick et al, 2008a). Field sampling campaigns can be directed and comprehensive, but are restricted to a few sites at some discrete snapshots in time, are weather limited, and affected by bloom “patchiness” (Foster et al, 2017). Continuous monitoring systems are constrained to a single (or a few) location(s) and miss spatial structure (Chang and Dickey, 2008). Remote sensing methods have the ability to image larger areas, but are limited in temporal resolution and spatial coverage is extremely sensitive to environmental conditions (eg. cloud coverage and glint) (Ruddick et al, 2008b; Urquhart et al, 2017). Yet, it is necessary to capture HAB temporal dynamics over hourly (e.g. diurnal variability, hydrodynamics, storms) to seasonal scales (growth/senescence cycles), and HAB spatial heterogeneity over scales *<*100 m (for predictive model parameterization and fundamental HAB scientific understanding) to *>*10 km (e.g. for tracking large scale historical intensity patterns). Beyond being useful for mitigation decision making, this information is also necessary to constrain these background HAB and environmental “signals” in order to better understand how HABs are affected due to anthropogenic or climatic effects. Capturing and measuring this variability is important to understand in situ HAB formation (Wells et al, 2015) and correctly quantify biomass system-wise.

Lake Okeechobee is the largest freshwater lake in Florida, with reports over the past 30 years of increasing HAB occurrence dominated by cyanobacteria (Betts et al, 2020; Wachnicka et al, 2022). In terms of temporal trends, while the public and scientific communities are well aware of an annual HAB season (Phlips et al, 2020), knowledge on local fast bloom dynamics is mostly anecdotal given its relatively rural location and until recently, few continuous water quality monitoring installations. As of 2021, the local water management agency South Florida Water Management District (SFWMD) maintains continuous monitoring sondes at several hot-spot locations with fluorometric biological and water quality sensors. Generally, in situ observations and sometimes remote sensing images suggest that these blooms can form, intensify, expand, displace, and disappear in a matter of a few days, but the exact scales of these motions are unknown – particularly for example, if blooms are expanding or simply moving. Furthermore, discrete snapshots from stations monitored bi-weekly by SFWMD show that these blooms might develop over various discrete – albeit repeated – regions in the lake (Wachnicka et al, 2022). Satellites on the other hand are constrained to, at maximum, daily captures of Lake Okeechobee, and this only when cloud coverage allows good visibility (which is unfortunately less frequent during bloom season). Furthermore, remote sensing product derivation is challenging due to the highly turbid environment (Urquhart et al, 2017). Thus, an overall paucity of HAB-related data over smaller temporal and spatial scales exists for Lake Okeechobee, especially when compared to other regions that have a longer history of operational in situ monitoring (e.g. Lake Erie) (Stauffer et al, 2019, and refs therein).

Autonomous mobile platforms offer an alternative solution to monitoring, covering a broad spatial range at a high temporal resolution (Beckler et al, 2019; Dunbabin et al, 2009; Hitz et al, 2012). The dual propulsion Nav2 Sail and Solar Drone (Navocean Inc.) is an Autonomous Surface Vehicle (ASV) that has been previously used for HAB detection and mapping during three *Karenia brevis* bloom events in the south-west Florida shelf between 2017 and 2018 (Beckler et al, 2019). Here, we describe one year of operations of a modified Nav2 equipped to monitor HAB and HAB-related conditions as part of a comprehensive, State of Florida-funded Lake Okeechobee monitoring effort (Harmful Algal Bloom Assessment of Lake Okeechobee program; HALO). The Nav2 patrolled around the clock (except for short maintenance periods) in the northern region of the lake, which is anecdotally considered an important area for bloom intensification. The Nav2 was equipped with fluorometric (chlorophyll-a, “chl-a”; phycocyanin, “phyco”; Colored-Dissolved Organic Matter, “CDOM”), suspended particulates (Red-Green-Blue backscatter), environmental (water temperature, conductivity and dissolved oxygen; DO) and meteorological sensors, as well as an Acoustic Doppler Current Profiler (ADCP; for 10 bins of current resolution in the vertical). The platform transmitted near real time conditions via cellular signal every 15 minutes (a set of scientific and navigation data), but also stored more than 2 million multivariable data points at high temporal frequency and small spatial scale. Here we describe vessel design, operations, and provide the first report of the long term results. We focus in particular on the fluorometric dataset given its purpose of providing a proxy for HAB presence. We review advantages and challenges of using this technology for HAB monitoring in a shallow turbid environments, and highlight major findings from the generated high-resolution unstructured dataset.

## 2 Materials and Methods

### 2.1 Nav2 Sailboat setup and operation

#### 2.1.1 Vessel design

The base design of the Nav2 Sail and Solar ASV (Navocean, Inc.) is detailed in Beckler et al (2019), including main body specifications, communication and navigation hardware, and energy systems. Vela is 2 m long vessel, weighting approximately 45 kg, and navigates at a slow steady speed for inherently safe operations. Vela was programmed for intelligent waypoint navigation under a broad range of weather conditions and is powered by renewable energy. Sensor payloads (and sensor “IDs” referenced throughout this text) are described in Table 1, with geolocated data collected near-surface (immediately below the boat at 0.15 m depth) at 30 second intervals. Meteorological measurements were taken 2 m above the surface. ADCP 3D current measurements were taken at 0.6 m depth interval (bins) throughout the water column to full lake depth, but discussion of ADCP data is beyond the scope of this work.

**Table 1.** ASV-mounted sensors and instruments.

| ID | Sensor | Manufact. | Parameter | Units | Range | Accuracy |
| --- | --- | --- | --- | --- | --- | --- |
| BB3 | Eco Puck | Seabird | Scatter 470 nm | m<sup>-1</sup> | 0 to 5 | 0.003 |
|  |  |  | " 530 nm | m<sup>-1</sup> | 0 to 5 | 0.003 |
|  |  |  | " 650 nm | m<sup>-1</sup> | 0 to 5 | 0.003 |
| CI | Cyclops Integrator | Turner Design | CDOM | ppb | 0 to 1500 | 0.15 |
|  |  |  | Chlorophyll | μg/l | 0 to 500 | 0.5 |
|  |  |  | Phycocyanin | ppb | 0 to 4500 | 2 |
| O2 | Oxygen Optode 4330F | Aanderaa | DO concentr. | ppb | 0 to 1000 | 2 |
|  |  |  | Air saturation | % | 0 to 300 | 1.5 |
|  |  |  | Temperature | °C | -5 to 40 | 0.03 |
| CT | Conduct. Sensor 4319B | Aanderaa | Conductivity | mS/cm | 0 to 75 | 0.018 |
|  |  |  | Temperature | °C | -5 to 40 | 0.03 |
| ATM | 200WX-IPX7 UWS | Airmar | Wind speed | m/s | 0 to 40 | 5% <sup>1</sup> |
|  |  |  | Wind direction | ° N | 0 to 359.9 | 3% <sup>1</sup> |
|  |  |  | Air Temperature | C° | -40 to 80 | 1.1 |
|  |  |  | Barom. Pressure | hPa | 300 to 1100 | 0.1 |
| GPS | - | - | Time | s | - | 1 |
|  |  |  | Lon/Lat | dd.mm | - | 10 m |
| CR6 | Data logger | Campbell Scientific | - | - | - | - |

<sup>1</sup>at 10 m/s winds

Sensor integration included custom housing design and machining and integration of a data logger (Campbell Scientific CR6) with customized cables and connectors. The meteorological sensor (Airmar) is a previously integrated component fundamental to ASV operations. Assembly also included software algorithm development and engineering to power and duty cycle the sensors, log the raw sensor data in the onboard data logger, and process the incoming data stream for sending over the Nav2 ASV near-real-time communications systems (cellular and satellite). The science water quality and ADCP sensors were set for one measurement/second with 5 seconds on and 25 seconds off for the BB3, CI, DO, and CT sensors.

Near-real-time data samples were compressed and transmitted every 15 minutes or more via satellite and cellular communications to a private server, and made publicly available through HALO web portal managed by the Gulf of Mexico Coastal Ocean Observing System (GCOOS) during operations for near-real time decision making. The ASV also recorded a complete data set to onboard memory, which formed the basis of the data analysis that is presented herein.

#### 2.1.2 Navigation

Operations started on December 31st, 2020, and navigation was controlled remotely by adjusting waypoints (Fig.2). The main objective was to repeatedly survey a polygonal area in NW Lake Okeechobee, encompassing multiple sites routinely monitored by SFWMD. Opportunistic, event-response redirection to emerging areas of interest was periodically performed, however, to conduct raster pattern surveys at spatial and temporal scales necessary to analyze HAB hot spots or other areas of interest as informed by other data collected as part of the HALO project, external reports, or per request of environmental managers.

**Fig. 1.**
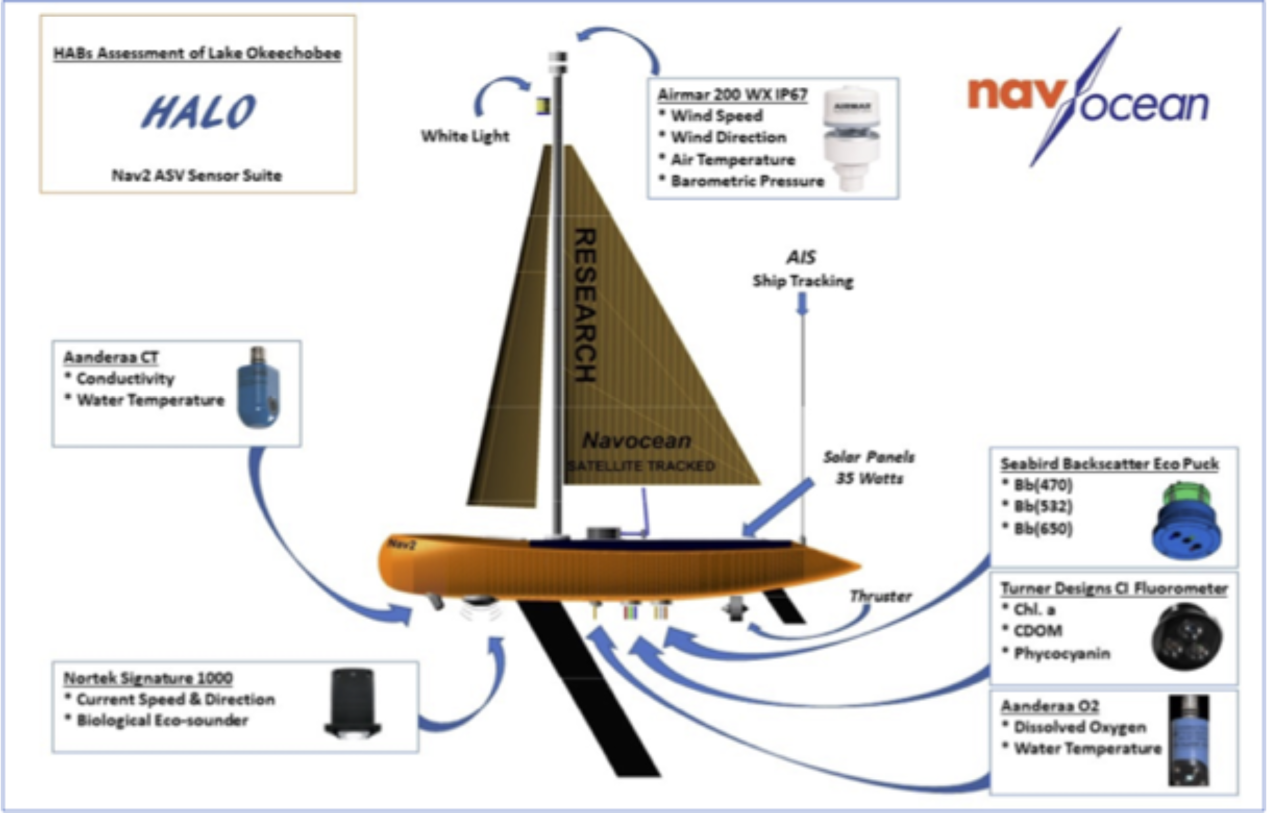
Nav2 Sail and Solar ASV showing the full sensor suite integration plan for this project.

**Fig. 2.**
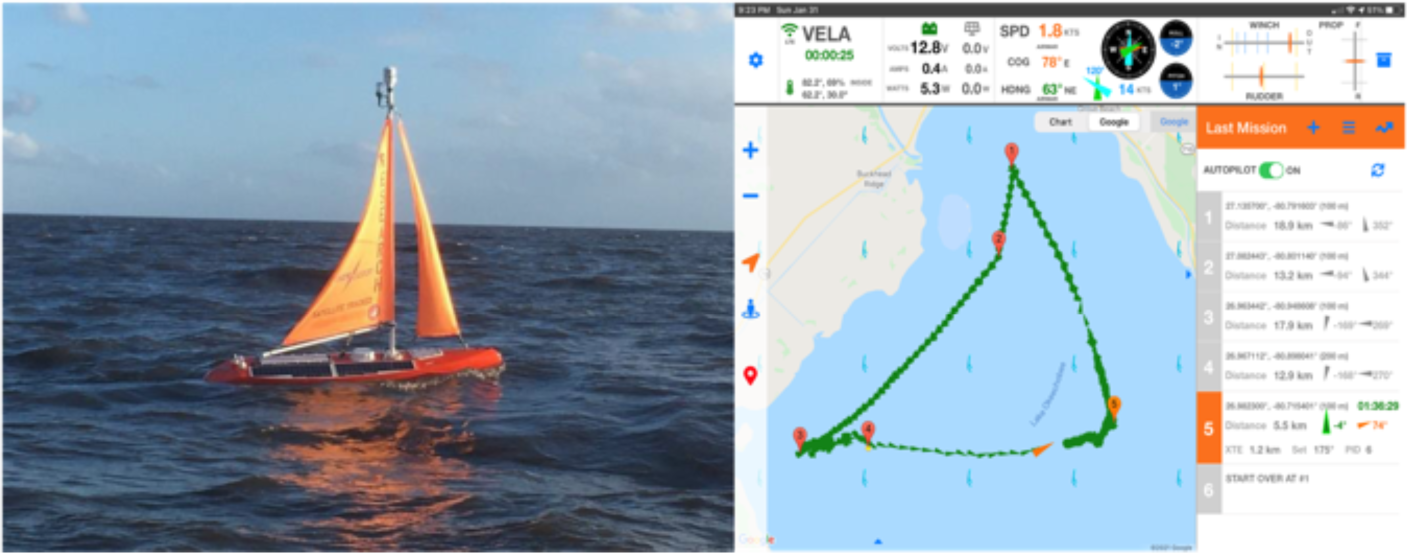
Vela launch on Dec 31st, 2020, sailing towards the L001 waypoint of the HALO Loop (left). Right: Mission Ops screenshots show Vela completing a HALO loop (right).

The main path, named the HALO loop, covered a region with frequent HAB occurance, and the semi - triangular path allowed for transect legs to be approached from the same direction every time. The HALO Loop was completed a total of 79 times until retrieval on January 1st, 2021. Some additional surveys were included as follows: 1) An algae hot-spot region in the north of the lake near the deployment site was discovered with elevated sensor readings confirmed visually and by water sampling. As such, a survey pattern called the “Taylor Creek Grid” (TC) was established in March and was surveyed opportunistically throughout the rest of the year at times when the Nav2 was returning to or launched from Taylor Creek before/after service. It is worth noting that this early detection of a cyanoHAB species (*Dolichospermum* sp.) was to our knowledge the first HAB reported in 2021, prior to detection via any other methods. 2) Vela was directed to sail from Taylor Creek on two side missions, one to Pahokee Marina (PM) in the southeast of the lake during a strong *Microcystis* bloom and another to the south/southwest end of the lake near Clewiston, to determine the feasibility of operating in other (shalower) areas of the lake and to survey a suspected *Microcystis* bloom in that area.

#### 2.1.3 Calibration and Validation

Prior to the first deployment, the CT and DO sensors were calibrated in the laboratory prior to the original deployment using their respective manufacturer-recommended procedure and sensor operating system, and the sensors directly reported these calibrated values during operations. Conductivity was calibrated using KCl solutions and temperature with a laboratory thermocouple. The DO sensor was calibrated using air-saturated laboratory solutions of deionized water at a known temperature and barometric pressure.

The fluorometric CI sensor and BB3 backscatter sensors reported RFU during operation, and individual channels were converted to desired units: chl-a and phyco in *µ*g/L, CDOM in units of *µ*g/L QSD, and RGB backscatter in turbidity units of NTU during data transmission and/or data processing. The BB3 used the initial factory calibration provided by Seabird to convert channel signal intensity to NTU for the duration of operations (true backscatter units can be derived in the future based on a simple conversion factor). Initially, the CI calibration response factors to convert RFU to desired units proceeded as follows: 1) chl-a was calibrated directly using a Chlorophyll-a solution (Turner designs) in acetone in a 5 cm quartz cell placed horizontally at the sensor head; 2) CDOM was calibrated using a quinine sulfate dihydrate (QSD) solution (Fisher Scientific) in deionized water; and 3) The phyco response factor was determine using a solution of Phycocyanin standard (Sigma-Aldrich) dissolved in water and held to the sensor head in in the 5 cm quartz cell, with the true standard concentration of the standard measured independently on a spectrophotometer to yield the corresponding concentration (Billy et al, 2023).

During servicing every 1-2 weeks, the BB3 and and CI sensors underwent check calibrations using a solid-state acrylic purple plexiglass sheet, similar to the methodology described in Earp et al (2011), in a custom fit housing ending (Fig.3). The housing was manufactured with different ring widths to create efficient and constant single-point calibration signal at a constant distance (0.375” for the BB3 R/G/B channels and CI CDOM channel, and 0.75” for the CI chl-a / phyco channels) sensors. Although calibrations for optical sensors are often only performed annually, a means to check for any variability due to biofouling or temperature effects was desired given that the sensors were to be deployed year-round. Visually, however, the instrument optical and housing surfaces remained free from biofouling growth in January through March and growth was only moderate during the warmer summer months.

**Fig. 3.**
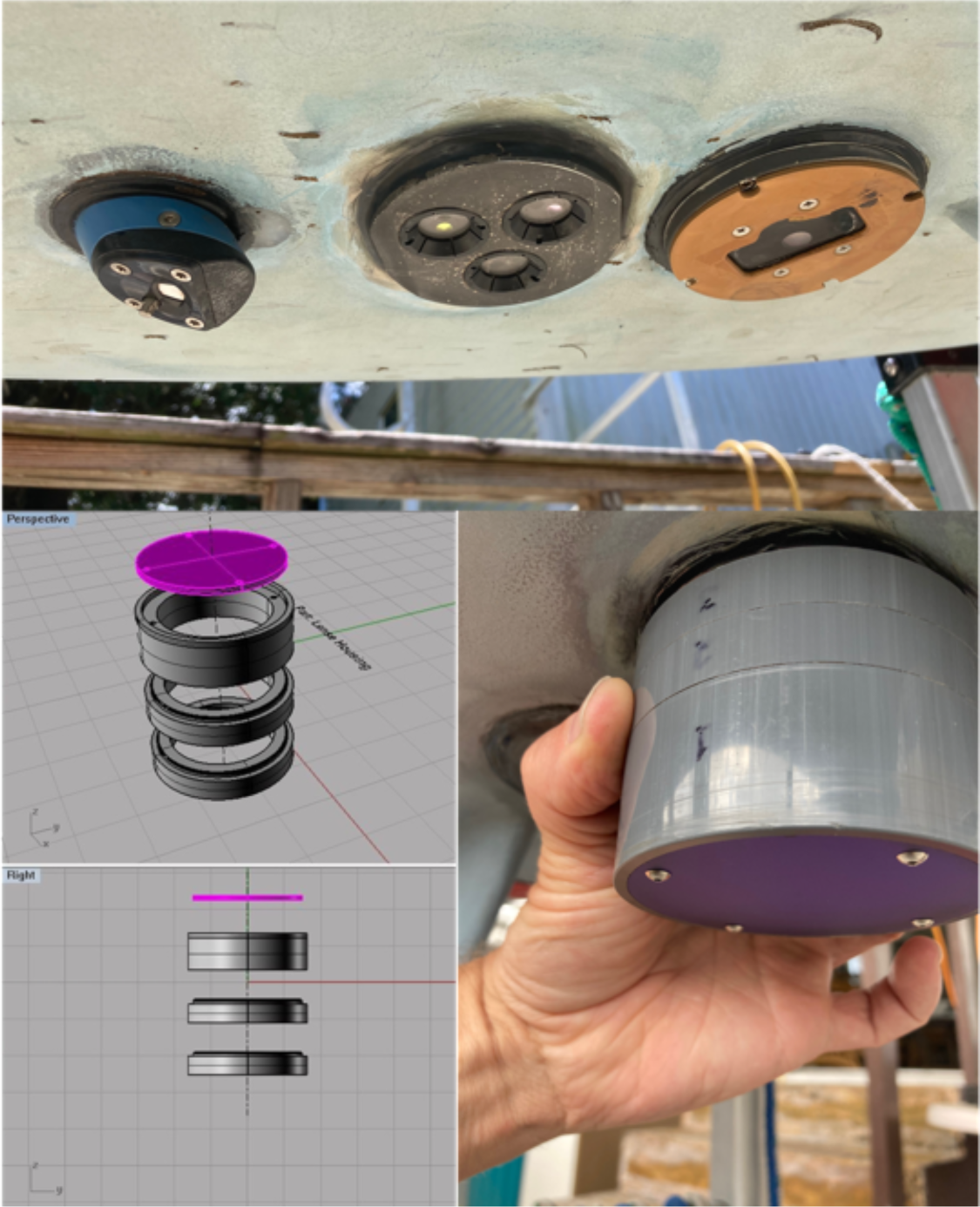
Custom validation cup developed to allow exposure of the CI and BB3 sensors to the SatinIce plexiglass sheet at discrete distances.

When logistically feasible, manual water grab validation samples were collected from the depth of the ASV sensors when the ASV was within 1 km of L001. Other validation samples were taken on side missions. All available values where used to correlate chl-a and phyco observations. Samples where analyzed for chlorophyll concentration and cell counts, as well as dominant taxa.

## 3 Results

The path traversed by Vela during the entire operation is shown in Fig.4. The HALO loop roughly connected several SFWMD monitoring sites marked in the map. Initial deployments lasted only a few days, as sensors required adjustments to reduce noise on turbid waters, but improvement allowed for up to two weeks of continuous operation over time. Data was collected from a total of 22 logger recoveries. For analyses purposes several geofences were established as follows: Right Track (RT), Bottom Track (BT), Left Track (LT), HALO Loop (loop), and Taylor Creek Grid (TC). The transect geofences (RT, BT, LT) were constructed to be around 1 km width. Results for Pahokee Marina (PM) and Taylor Creek (TC) side missions are presented in a separate section.

**Fig. 4.**
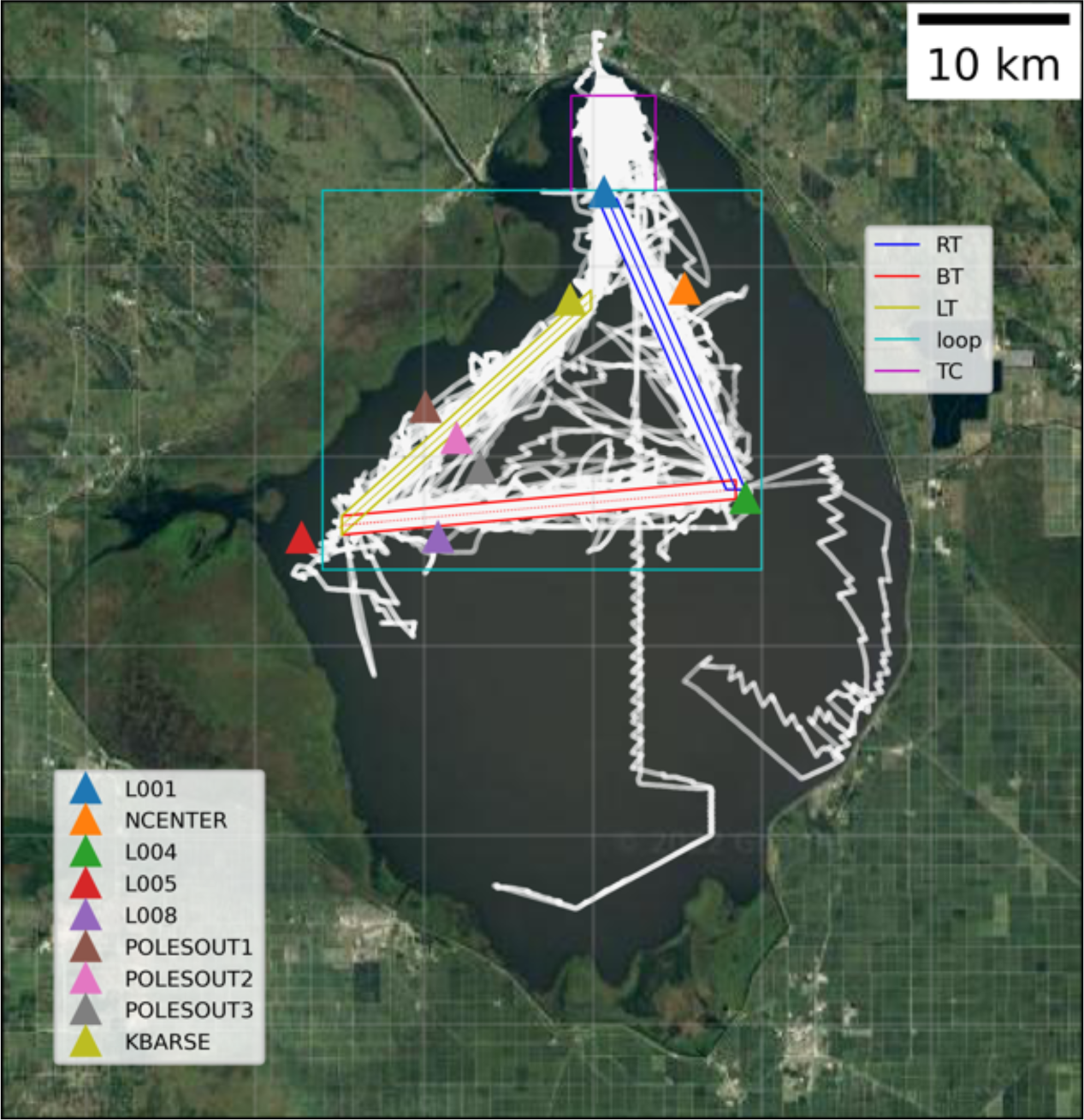
Path traced by Nav2 during 2021 in Lake Okeechobee(white). Colored polygons show data geofences for the right transect (RT), bottom transect (BT), left transect (LT), HALO loop (loop) and Taylor Creek Grid (TC). Side missions to Clewiston and the Pahokee Marina area (PM) are visible towards the south and southeast of the lake, respectively. Markers show the positions of relevant stations monitored regularly by the SFWMD.

Quality control included the use of a range limit (allowable min and max), and removal of spikes (unaccounted noise) defined as a jump in consecutive measurements larger than a threshold. These spikes are most likely caused due to sudden obstruction of sensors by debris during navigation or electrical noise.

After this cleanup, more than 4.3 million quality-controlled data points were collected for each sensor. Validations of these calibrated values are discussed later.

### 3.1 Temporal trends

#### 3.1.1 Seasonality

The full year of mobile time-series constrained to the HALO loop geofence for each water variable is shown in Fig.5. This time-series is filtered to a 30 seconds mean, and a daily mean is superimposed.

**Fig. 5.**
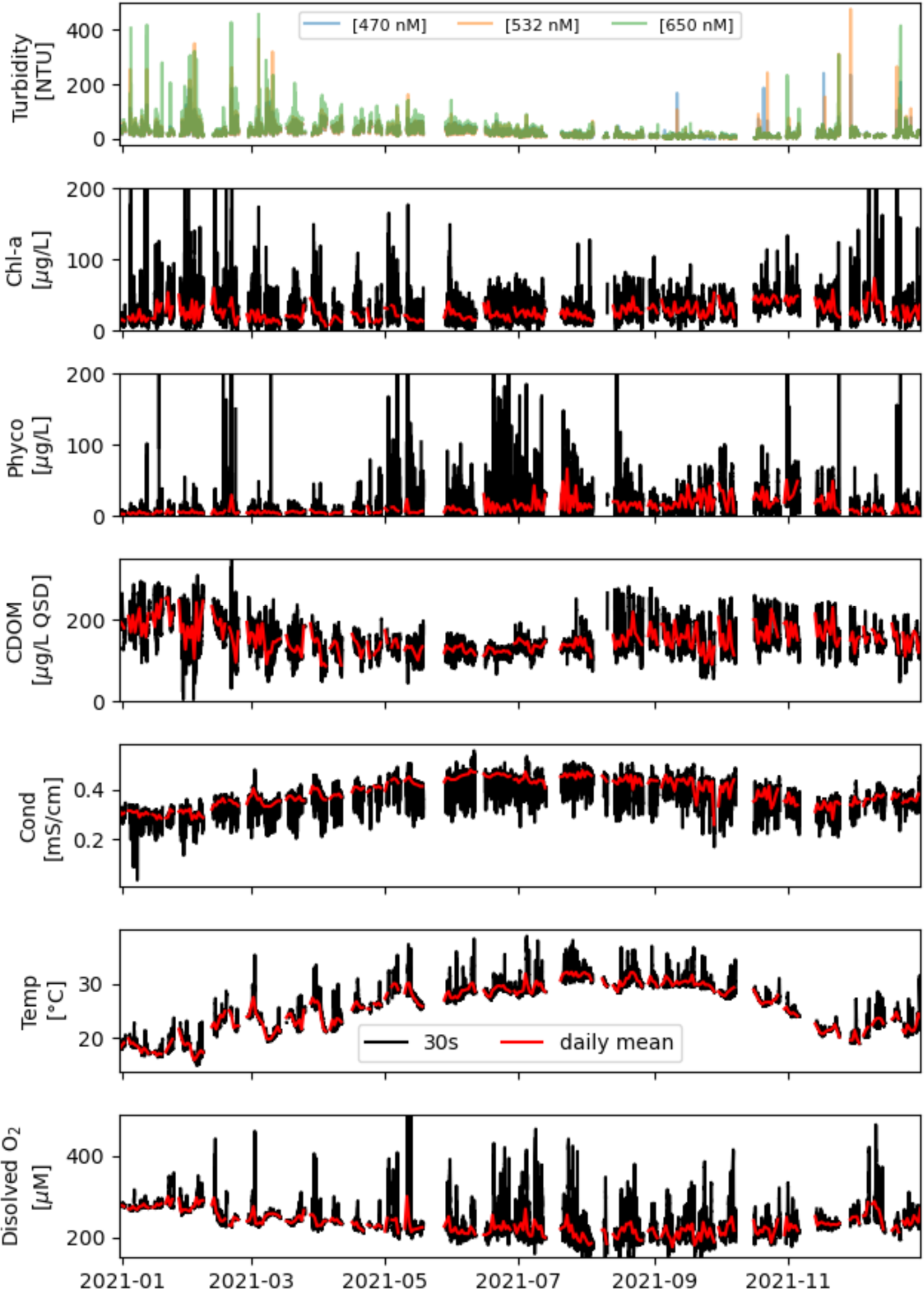
Mobile time series of surface water properties recorded by the Nav2 within the HALO Loop geofence.

Higher turbidities and variability in chlorophyll were recorded during winter and spring months (beginning and end of the year). Isolated extreme peaks in phyco where recorded as early as mid-February and early May, associated with a relatively high chl-a signal. Presence of algal patches in these instances were confirmed on site, with the dominant taxa being *Dolichospermum sp*. Consistently higher signals in phyco where recorded between mid June and late November (i.e. daily means *>*10 *µ*g/L), but with a relatively lower chl-a signal. This seasonal peak coincided with *Microcystis* dominance, corroborated from water samples collected during this period (not shown). CDOM, while highly variable, had a trend inverse to conductivity (r = −0.51), with consistently lower values between April and mid August (i.e. daily mean *<*150 *µ*g/L QSU). The full seasonal signal was captured by other environmental measurements, with overall higher temperature and conductivity, and lower DO in summer months.

#### 3.1.2 Diurnal and higher frequency variability

High frequency variability is difficult to observe in the full dataset (Fig.5), but can be contrasted between the raw (30 sec mean) and the daily mean signal. In general, a strong diurnal signal was captured in the environmental parameters (temperature, conductivity, and DO) but this is not always the case, for example in early year missions under strong wind conditions.

### 3.2 Large-scale spatial variability

The presence of patterns and variability is also strong spatially. To analyze the spatial trends, data points within the transect geofences (LT, RT and BT) where projected to the center line. To illustrate the captured motions the time-space plane of the left transect (LT), also termed coastal transect, is shown in Fig.6. RT and BT planes are presented in the supplement as Fig.A1, and not further discussed. In the LT plane the improvement in the quality of operations is visible in the more continuous visualization beginning in May. Missions lasted longer and navigation accuracy was improved over time, by adjusting sailing patterns. This accuracy improved the density of data falling in the transect geofences, permitting better space-time pattern reconstruction.

**Fig. 6.**
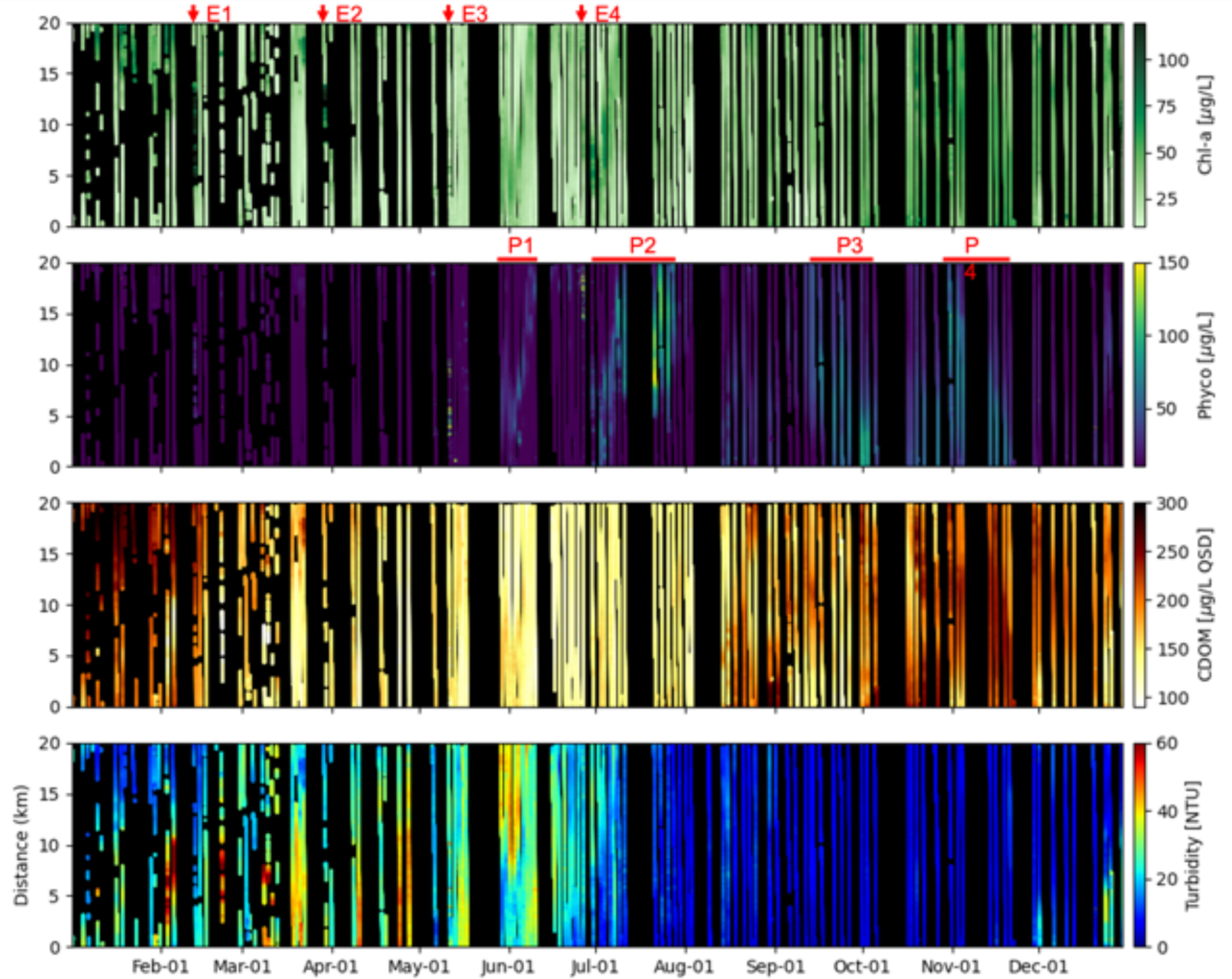
Spatio-temporal plane depicting optical property characteristics along LT (see Fig.4). The vertical axis shows the projected distance along the transect, starting from the SW end. Red markings indicate short bloom events (E 1 to 4) and longer periods of bloom motion and growth (P 1 to 4) discussed in the text.

The first short duration episodes HAB’s are visible as isolated spots appearing between February and April (mostly in the middle of the transect), characterized by high chl-a and slightly elevated phyco (E1 and E2). Dominant species extracted from water samples collected on the site were mainly *Dolichospermum*, with some presence of *Microcystis* in the later episodes. An event in May, with greater phyco relative to chl-a, is the first appearance of a *Microcystis* bloom (E3). Later, in June, a continuous record of elevated phyco indicates the motion of a bloom in the NE direction along the coast, revealed by a sloping of the signal as represented on the spatiotemporal plane (P1). While not an extreme bloom, P1 is the most continuous record of bloom motion in our data, showing a relatively constant displacement speed of aproximately 1.5km per day (close to 18 km in 12 days). Relatively high but short-term blooms continue to be active in the northern end in June, with a clear high phyco concentration peak occurring by the end of the month (E4). In July the *Microcystis* bloom formed at the SW end, was transported over time towards the center of the transect, and expanded to occupy the upper 15 km of the transect (P2). While the first sloping section in the elevated phyco and chl-a signals indicate motion, the second half indicates radial expansion or growth. This bloom persisted approximately one month. Higher concentrations of phyco remained throughout the rest of the year from August onward, but two more extreme growth and displacement events are visible in the end of October (P3) and end of November (P4).The slope of these two events are reversed, indicating a SW motion.

### 3.3 HAB motion detection example

The northeastern HAB motion P1 identified in Fig.6, starting in late May and captured with high resolution in a 2-week record in June is discussed here. Sentinel 2 images show highly turbid waters, and dark green coloration at the south end of the transect on 05/25/2021. Roughly two weeks later, on 06/29/2021, a cyanobacterial bloom is fully developed in the region, following the coastline. This indicates that the bloom started forming some time after 05/25/2021. The development of this event is captured by a total of 9 transects covered between 05/29/2021 and 06/10/2021, roughly every 1.5 days (Fig.7). When the Nav2 starts the continuous mission on 5/29/21 after servicing, a moderate HAB hotspot is located at the SW portion of LT. Because the Nav2 operates 24/7, day/night conditions should be considered as factors possibly impacting the measurements. Tracks mostly in the daylight (1,3,5,8) show consistently higher Chl-a values than night tracks probably due to the sensor sensitivity to light, but this is not the case for phyco. No major displacement is observed in the chl-a until tracks 3 to 5, when the center of the hotspot is displaced northwards about 3 km (in 3 days). During track 5, phyco concentrations are divided in two hotspots, separated about 5 km. Over the last days (tracks 7,8 and 9) the HAB hotspot has moved toward the NE end of the transect. Some additional observations can be made from these records. Fully night-time tracks show lower chl-a, due to probably the sensor sensitivity to light. This effect however is not observed in the phyco, where peaks are as elevated or higher than the day tracks. Phyco and chl-a peaks are not co-located, suggesting different population compositions of the blooms, but the overall bloom motion can be followed from the SW toward the NE end of the transect over the two weeks records.

**Fig. 7.**
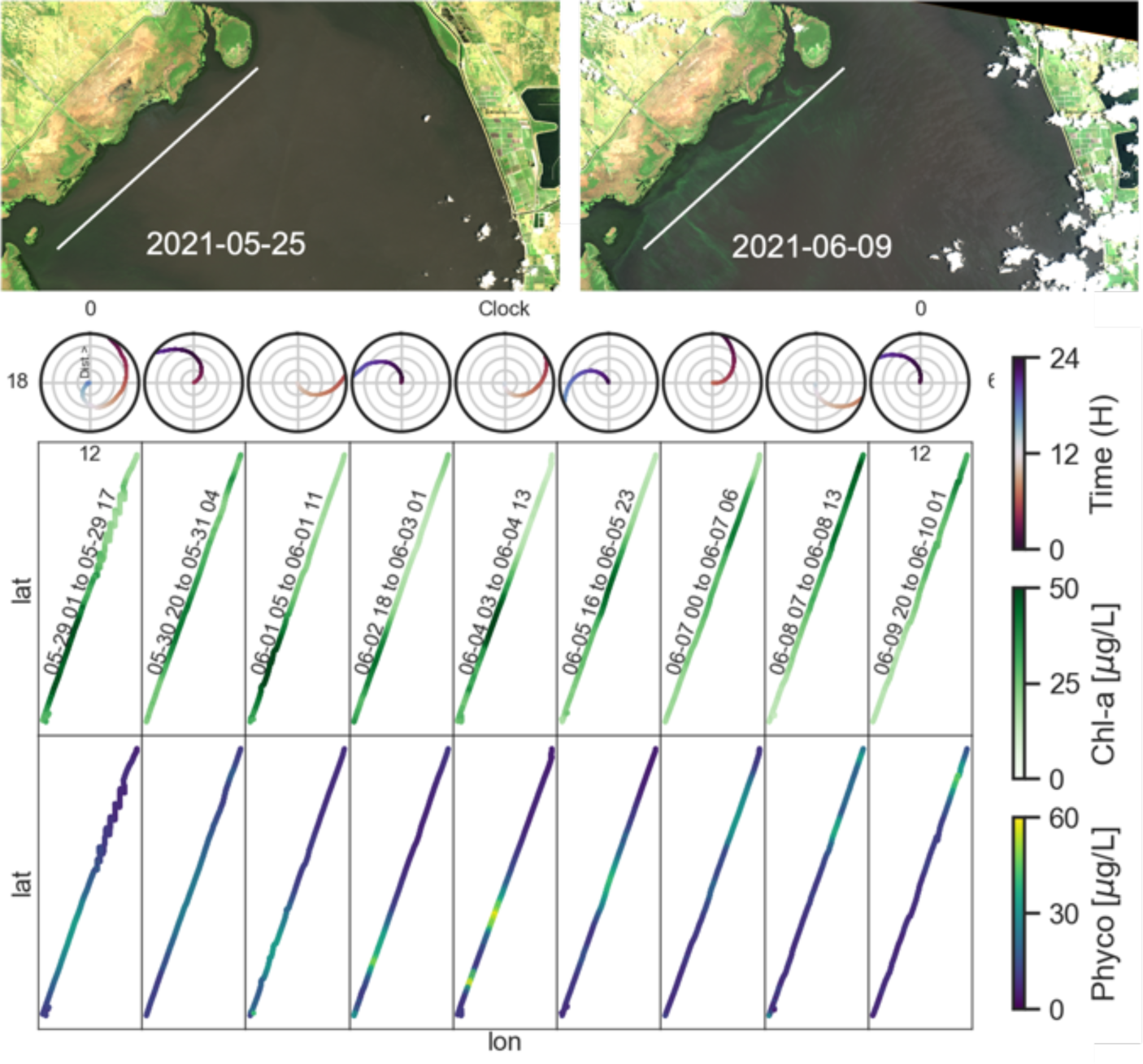
Bloom detection (P1 in Fig.6). In descending rows: 1) Sentinel 2 true color processed images for 05/25/21 and 06/09./21 These are the only clear images of the region close to the bloom displacement event captured between 05/29/21 and 06/10/21. 2) Time of the day indication of the 9 tracks that captured the event. The radial distance is the track distance from the SW extreme, and the angle (and color) the time in decimal hours. Light colored tracks in the lower half of the clock are in daylight (6-18 H), and the dark colored in the upper half is night-time. 3) Chl-a and 4) Phyco captured in lon/lat space for the 9 tracks.

### 3.4 HAB fingerprinting using spatial structure

Below, we demonstrate how high-resolution fluorometer spatial variability can potentially be used as a tracer of bloom compositions, a useful tool potentially not reproducible by any other sensing platform type. Analyzing the full data set, we found a complex relationship between the relative fluorescence of chl-a and phyco as shown in Fig.8. Here we use direct sensor output RFU instead of converted concentration values, given the relatively low confidence of the calibrated values and the anticipated use of this same sensor payload (i.e. the same Turner CI for both chl-a and phyco) in this same lake environment in the future. The scatter plot shows two distinct correlation lines (i.e. above and below the 1:1 line respectively) that bound a grouping of intermediate points. Given this evidence, we suspect that the ratio between phyco and chl-a may be a useful indicator to distinguish and identify dominant phytoplankton species assemblages (i.e. along either of the two “slopes”), whereas a more mixed-assemblage may instead plot somewhere between these respective linear response regions. Various HAB or other phytoplankton species certainly do display variable phyco relative to chl fluorescence with the rationale that phyco is specific to cyanobacteria whereas chl-a is common to all phytoplankton (Foy, 1993; Rousso et al, 2022). For instance, data from the LT is generally more distributed in the center of the the two slopes than the RT or BT (8). Indeed, the LT is generally a biologically dynamic region with high phytoplankton species diversity, given relatively clear water and nutrient inputs from the Kissimmee River (northeast) and Fisheating Creek (west) (Welch et al, 2019; Wachnicka et al, 2022).

**Fig. 8.**
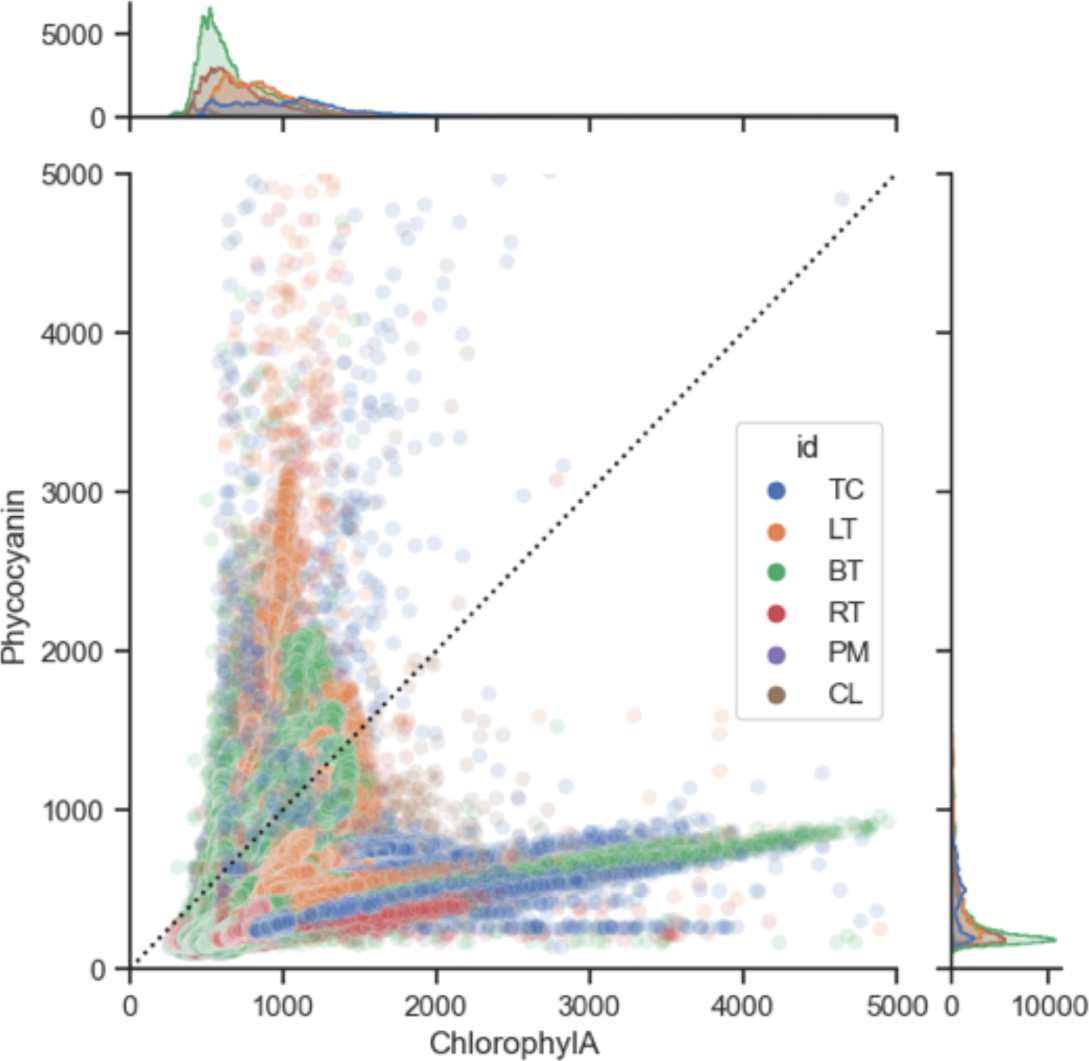
Chlorophyll-a and phyco relationship. The hue indicates the different regions as shown in Fig.4 (points inside the loop region not falling into transects have been removed for clarity).

Additional bloom fingerprinting capabilities may be gleaned when we consider the phyco/chl-a ratio expressed within the context of its spatial variability. We present two early-year side missions excerpts that use data directly output to the web portal, showing the true near real-time capabilities of the Nav2 for HAB early warning. The Taylor Creek bloom was surveyed on 4/21/21, and the Pahokee Marina bloom surveyed on 4/26/21. Fig.9 shows the location and detailed sail paths of a few hours of survey, displaying the phyco/chl-a ratio. While chl-a values were elevated in TC, the overall variability in the phyco/chl-a ratio was small (0.06 std) when compared to PM (3.75 std), a factor of 62.5x greater. This is visible in the frequency distribution for both tracks (inset in Fig.9), and also in the intense patchiness observed in the PM track.

**Fig. 9.**
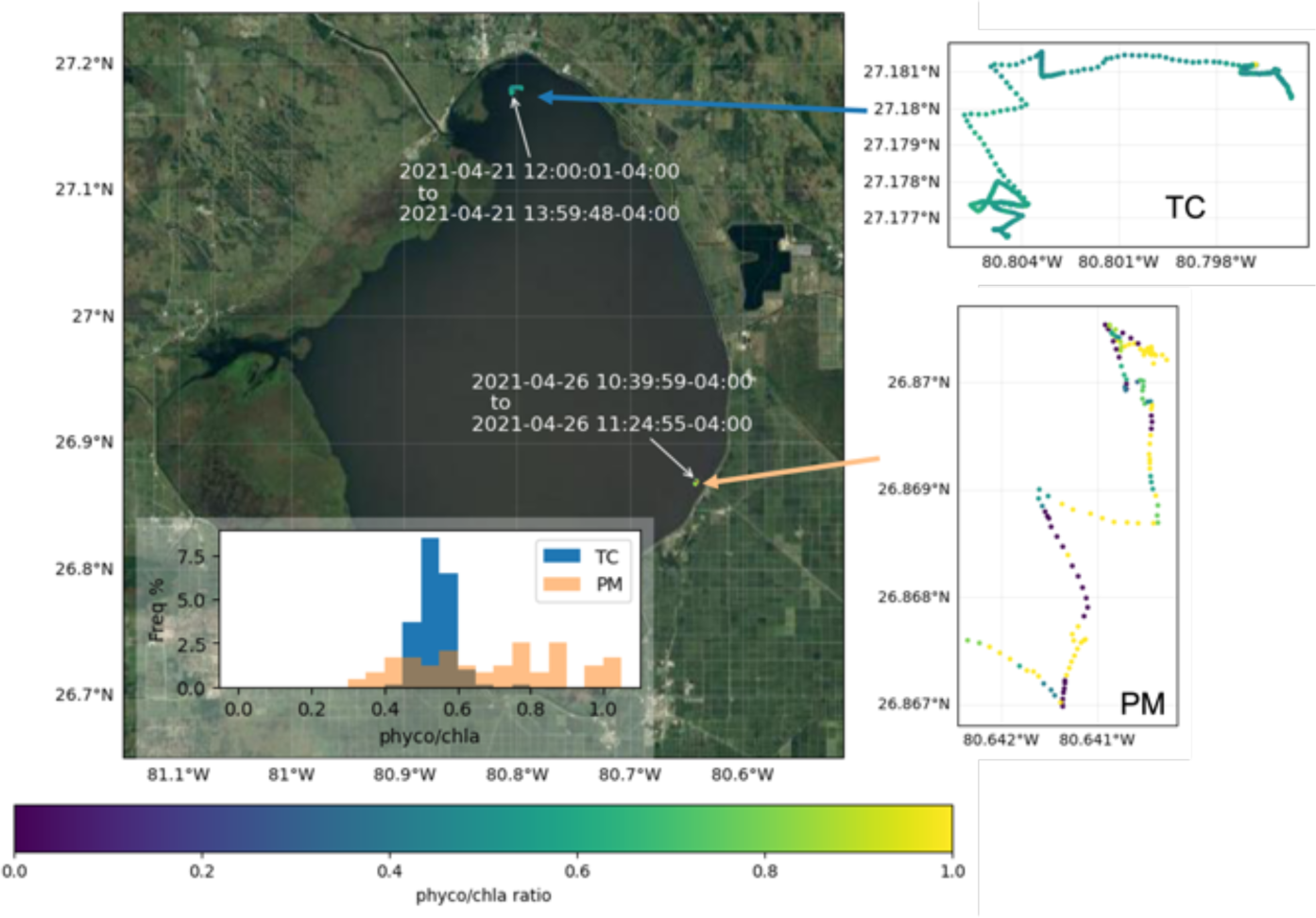
Near real-time acquired data showing ratios of phyco to chl-a (in RFU) for two segments into the Pahokee Marina (PM) and a Taylor Creek (TC) missions separated by 5 days. The individual sail paths are zoomed in to show the fine spatial variability. The normalized frequency distribution of the ratio is shown in the inset.

**Fig. 10.**
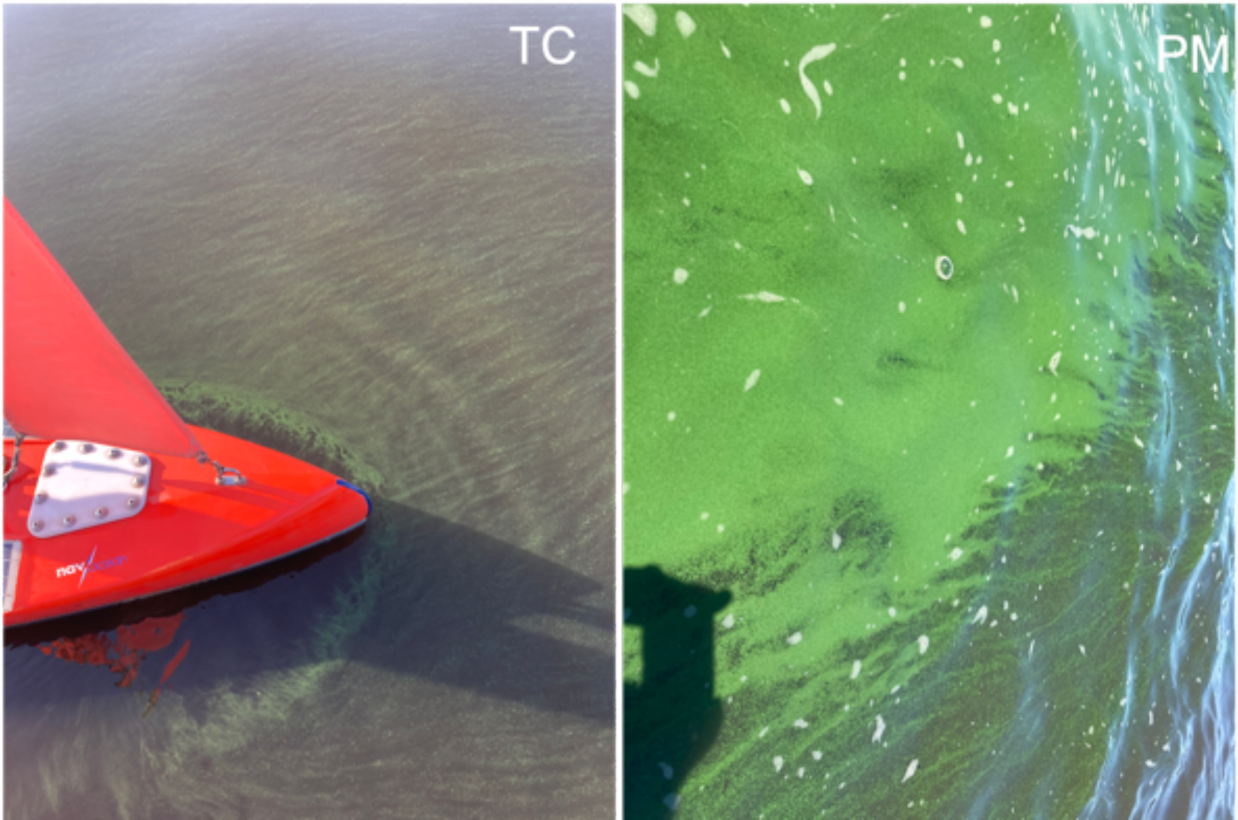
Photo images of the surface scums captured during a visual inspection of the operation in Fig.9, at Taylor Creek (TC) and Pahokee Marina (PM).

Visual inspections of the sites during the operation confirmed the presence of blooms in both sites, but with an obviously different surface scum appearance (see Fig.9). The individual signals of chl-a and phyco for both tracks are included in the supplementary material (Fig.A2), for which PM was more variable than TC by a factor of 7.6x for Chl-a and 60.7x for Phyco. Grab samples under the microscope showed that TC was dominated by *Dolichospermum*, another toxic cyanobacteria found in Lake Okeechobee, and *Microcystis* was present in a lower abundance with some of the colonies aggregated with *Dolichospermum*. At the surface the scum appeared as a “streaky” but relatively homogeneous deep green (Fig.9, left). In PM, both *Microcystis* and *Dolichospermum* were also present but *Microcystis* was the dominant taxa based on the size of the colonies. Visually, this bloom showed as a more patchy bright green layer with surface scums, with many marked edges as shown in Fig.9 (right). Thus, the ASV-acquired data may uniquely provide additional bloom assemblage discrimination capabilities based on the overall macro-structure (0.1 - 1 km range). This unique capability of the Nav2 is ripe for operationalization of a fingerprinting algorithm for HAB detection, although the introduction of both multivariable and spatial dimensions would require advanced model development.

**Fig. 11.**
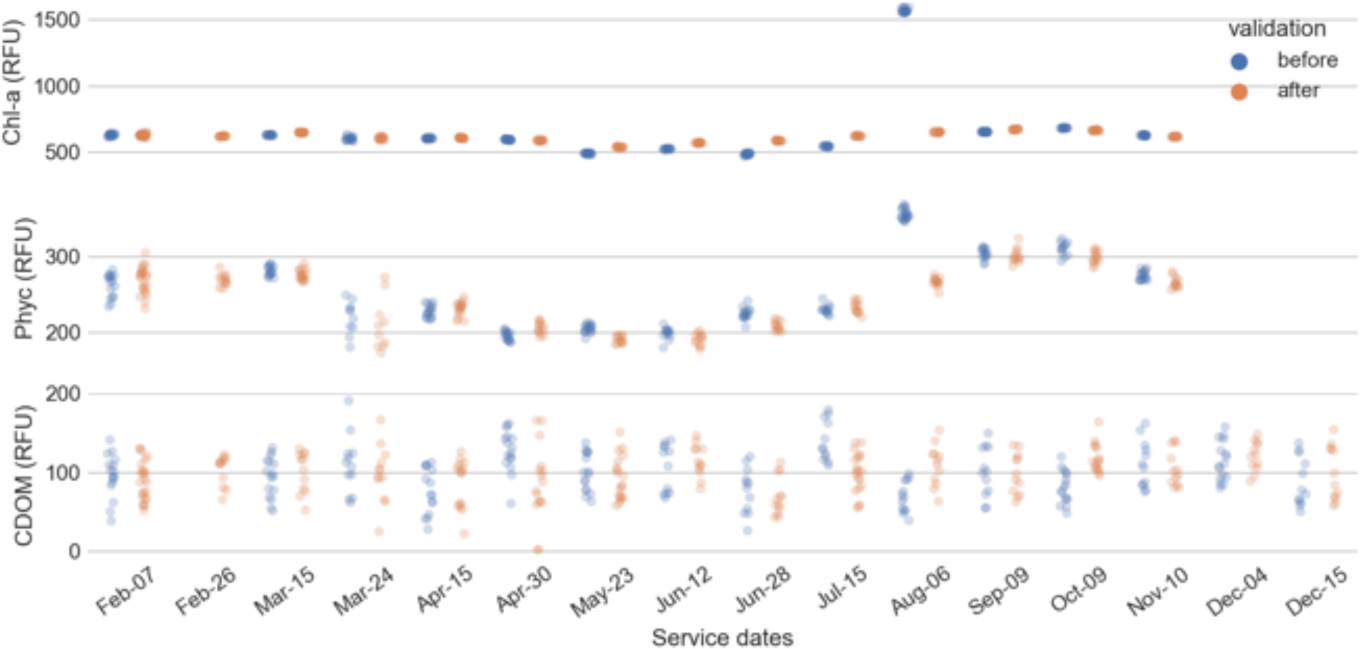
Solid-state validation checks for dark, and before and after cleaning of sensors during routine service.

### 3.5 Validation

Validation grab samples collected from the field and subject to laboratory measurement generally showed poor correlation with the in situ Nav2 measurements for chl-a (rms 182 *µ*g/L). We attribute this due to either to heterogeneity of grab samples versus in situ measurements, the high turbidity of the lake system (affecting fluorescence measurements), or the high CDOM (affecting backscatter measurements). However, through the continuous use of the sensors and clear indication of both short and long-term spatiotemporal patterns in the sensor signals, we are confident that the sensors are capturing real trends, especially within the context of the primary mission purpose of detecting HABs for rapid mitigation purposes. Solid-state optical sensor validation field checks provided a relatively constant baseline response for sensor performance between service dates (Fig. 6). Chl-a and phyco sensor performance was consistent with minimal variation before and after cleaning the sensors with the exception of the August service date. Removing this outlier, the fluorescence sensitivity loss for every service time t, defined as after [t] - before [t+1], was on average under 5% for chl-a and under 1% for phyco. Some seasonal fluctuation is also observed in the phyco checks, and we attribute this to possibly a temperature effect, especially given that validation was performed under air temperature conditions and the CI is not temperature compensated (e.g. Blaen et al, 2017). Solid-state field checks of CDOM readings were less consistent, with high variability within the before and after readings, and even within individual validation steps. This might indicate that the purple lens utilized in the validation is not appropriate for this type of sensor and some alternative methods should be investigated. Ultimately, however, the lack of intensity loss from biofouling and the stability of the checks over time give confidence to the relative temporal variability of fluorometer data.

## 4 Discussion

Lake Okeechobee (1732 km^2^ area) is sufficiently large to allow the development of complex dynamic features in its water column (Ji and Jin, 2006; Jin et al, 2002; Jin and Ji, 2004), environmental composition and in turn, biogeochemical conditions (Carrick et al, 1994; Stockwell et al, 2020) including HABs (Beaver et al, 2013). Wind velocity is responsible for the main surface current and modeling studies have shown that strong wind events can produce the complete mixing of the entire water column, while thermal stratification is present during calm weather periods (Jin et al, 2002). On the other hand, wind driven waves are responsible for sediment (and thus sediment nutrient load) resuspension, while wind induced currents transport those sediments (Jin and Sun, 2007). This heterogeneity is most likely responsible for the variability in the water quality and the frequency and composition of HABs in different regions of the lake as reported by SFWMD (Welch et al, 2019; Wachnicka et al, 2022),

Accordingly, SFWMD dramatically increased in Mar. 2020 their number of routine phytoplankton and water quality monitoring stations, and implemented the use of continuous-real-time surface and bottom monitoring sondes Wachnicka et al (2022).

Use of the Nav2 ASV during the HALO project complemented these routine monitoring efforts and allowed the identification and even discrimination of cyanoblooms very early in the season. While we identified typical, expected seasonal signals during our year long continuous monitoring campaign (Fig.5), we also identified that spatio-temporal variability can be several orders of magnitude larger than the seasonal differences (i.e. see phyco and chl-a panels in Fig.5). An accurate accounting for this variability is critical for both understanding climatic-scale trends but more importantly, for implementing an appropropriate HAB mitigation response strategy.

The magnitude of this variability needs to be investigated further at various spatio-temporal resolutions, and this massive, comprehensive in situ data-set will provide further insights into this question. While this work aims to be a first report on our findings, future work would include elucidating the differences in the variability as contrasted with routine monitoring. The HALO project additionally collected many other types of multi-platform data that will form the basis for a comparison.

Fixed-location monitoring installations give useful information on HAB and water quality evolution at large time scales (years to decades) and can be used to provide alerts when high concentrations of toxic algae are present in a station. However, even when using a considerable number of stations, this strategy will still not provide the ability to measure high-resolution spatial responses, the spread of a bloom event, or to identify localized features not covered by the station network. Satellite images on the other hand, can help identify lake wide distribution patterns of color properties, which are indicative of algal blooms, turbidity or CDOM, but the small-scale temporal variability and environmental conditions are not captured.

The Nav2 platform presents some advantages over these more conventional techniques. Operations directly provide high spatial and temporal resolution information on important biological, water quality, and environmental properties. Excitingly, the continuously transiting in-water vehicle appears to capture macro-scale bloom structure. We can envision a HAB species fingerprinting technique in which a near real-time algorithm uses a recent look-back period (i.e. travel distance) to examine fine spatial-scale spatial variability, e.g. the standard deviation of the phyco/chl-a ratio, to provide a HAB genus-level identification capability more specific than just indicating the present of a cyanobloom based on a green color (e.g. cyanobacterial index). The use of fluorecence phyco/chl-a ratios to discriminate species composition has been proved in culture experiments (Rousso et al, 2022) and more cleaner environments (Haggard et al, 2023). Implementing thresholds within a turbid ambient such as lake Okeechobee is a challenge, but our results are promising. This is critical given the cost of mobilizing mitigation response resources in response to a cyanobloom, despite a non-zero chance of this being a “false alarm” if the species of interest is non-toxic.

The sailboat has traversed over 10,000 km of the lake in a semi-triangular 72 km path, and additionally targeted detailed investigations of bloom events and side missions. A total of 6,888 hours of operation provided more than 4 million datapoints for every sensor used. Using the operational path described in this study with a single Nav2, we managed to survey three separate transect legs at a repeat frequency of between 1 and 3 days, depending on the wind.

For the specific period of time and transect shown in 7, specifically, a repeat frequency of approximately 1.5 days was obtained. This repeat frequency is likely sufficient for mobilizing HAB mitigation resources to ensure blooms are mitigated while still tractable (i.e. prior to expanding rapidly). There are major opportunities for improvement of autonomous path planning, particularly in response to anticipated/modeled wind conditions. Mission objectives will also highly affect repeat frequency; for example, a single transect could be surveyed in the forward and reverse directions, gaining a higher repeat visit frequency in the center of the track. This may be ideal for more targeted applications such as monitoring near water intake structures or a known contributing HAB source (e.g. Maumee Bay, Lake Erie) (Matson et al, 2020).

While extensive future work remains on this dataset, some highlights on this first report include: This was the first ever effort in Lake Okeechobee to measure CDOM and phyco in situ at large scale. The enormous number of data points collected allow differentiation of daily and sub-daily frequencies, as well as to cover seasonal trends and responses to specific events. We could also identify some combined spatiotemporal trends, and further analysis supplemented with satellite images and in situ measurements provide insight into spatial spread of bloom formation, development and decline. The measurements represent a big data challenge that can elucidate unexpected results. The data set is complex given the interwoven spatiotemporal dimensions, but modern statistical analysis tools and use of supercomputing power provide multiple approaches. This first report only highlights the potential of this type of platform for HAB monitoring, and we expect this dataset to generate many more insights from ongoing data processing.

## Supplementary information

In Appendix.

## Acknowledgments

This study was part of the Harmful Algal Bloom Assessment of Lake Okeechobee (HALO) Program, which was supported by the Florida Department of Environmental Protection (FDEP) Innovative Technologies for HAB Mitigation Grant MN016. The authors acknowledge other PI’s and researchers involved in the project including Dr. Jordon Beckler’s and Dr. Malcolm McFarland’s group members that assisted with field operations and field validation, especially Lynn Wilking and Jessica Carney for the performing of lab calibrations and water sample analysis for colonies, the GCOOS effort lead by Dr. Barb Kirkpatrick for creating and maintaining the HALO portal which provided near-real time updates during operations, Dr. Tim Moore for discussions relating to remote sensing products, the Navocean team for the deployment and operation of the Nav2, and several local residents of the town of Okeechobee (FL) for assistance with operations.

## Declarations

### Competing interests

S. Duncan is the Founder and President of Navocean Inc., the provider of the Nav2 Sail and Solar Drone.

### Availability of data and materials

The dataset generated during and/or analysed during the current study is available in a zenodo repository: https://doi.org/10.5281/zenodo.8264853.

## Appendix A Supplementary figures

**Fig. A1.**
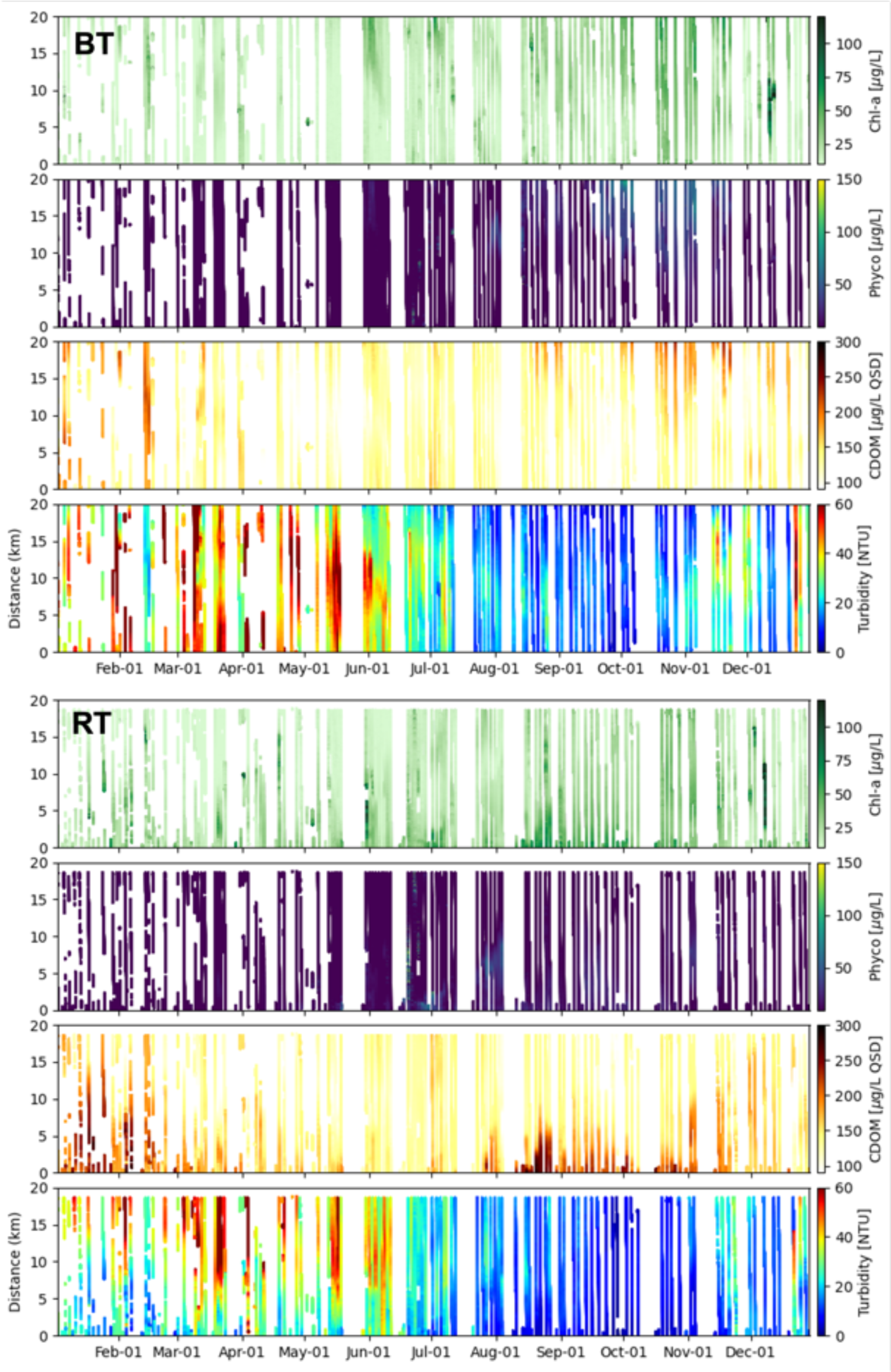
Spatio-temporal plane depicting optical measurements along RT and BT as defined in Fig.4), and comparable to Fig.6. The vertical axis shows the projected distance along the transect, starting from the north for RT, and west for BT.

**Fig. A2.**
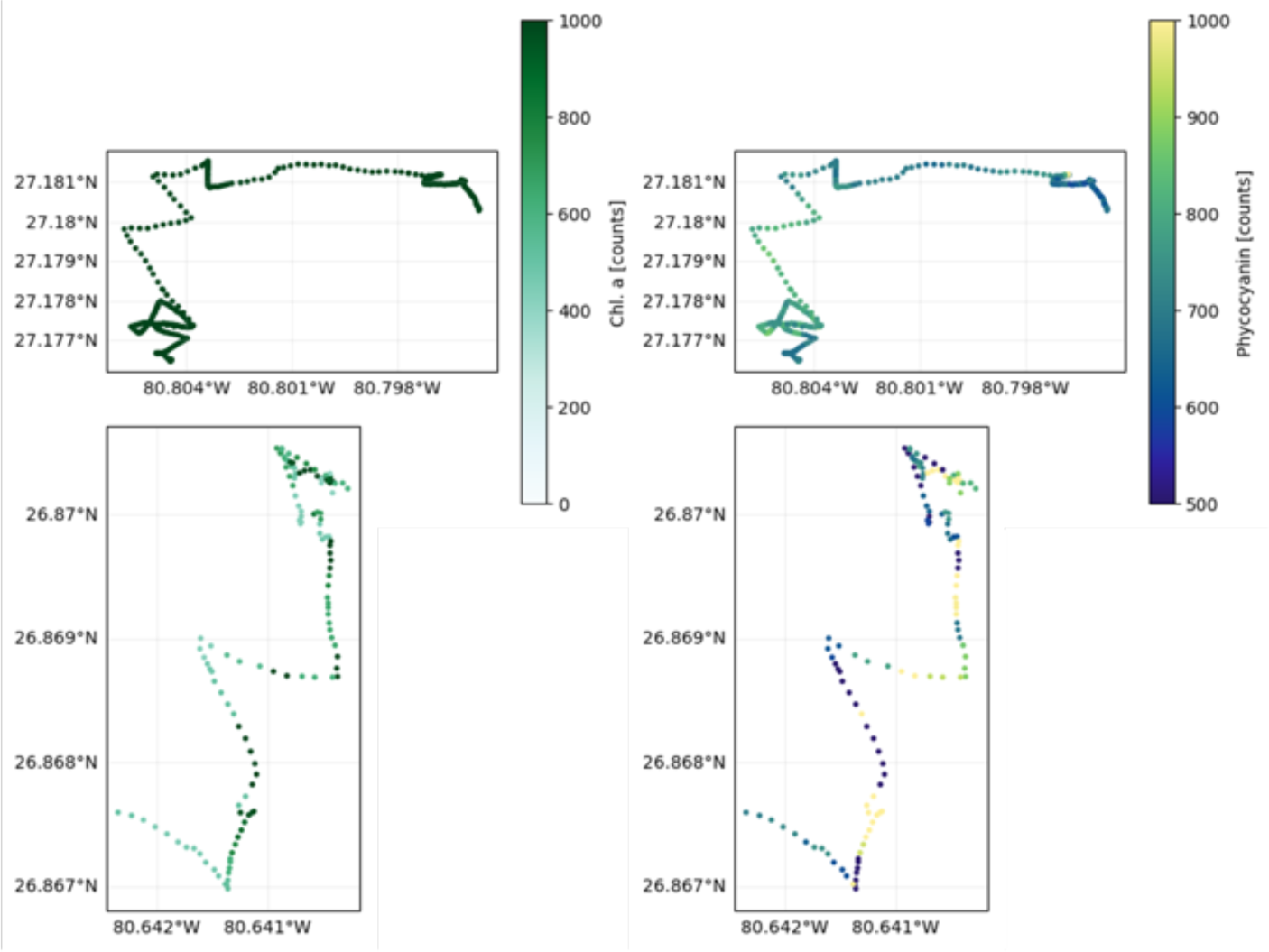
Nav2 paths as in Fig.9 showing chl-a and phyco (in RFU) for two segments into the Pahokee Marina (PM) and a Taylor Creek (TC) missions.

## Notes

https://doi.org/10.5281/zenodo.8264853

### Competing Interest Statement

This study focused on the performance of an autonomous sail-powered vehicle used for monitoring harmful algal blooms. Author Scott Duncan is the Founder and President of Navocean Inc., the provider of this Nav2 Sail and Solar Drone.

## References

Babin M, Roesler CS, Cullen JJ (2008) Real-time coastal observing systems for marine ecosystem dynamics and harmful algal blooms : theory, instrumentation and modelling

Beaver JR, Casamatta DA, East TL, et al (2013) Extreme weather events influence the phytoplankton community structure in a large lowland subtropical lake (Lake Okeechobee, Florida, USA). Hydrobiologia 709(1):213–226. 10.1007/s10750-013-1451-7

Beckler JS, Arutunian E, Moore T, et al (2019) Coastal Harmful Algae Bloom Monitoring via a Sustainable, Sail-Powered Mobile Platform. Frontiers in Marine Science 6:587. 10.3389/fmars.2019.00587

Betts A, Jones P, Ollis S, et al (2020) Appendix 8A-1 : Lake Okeechobee Watershed Protection Plan 2020 Update. Tech. rep., South Florida Water Management District, West Palm Beach, FL

Billy J, Pruvost J, Ĺepine O, et al (2023) Development of a spectrophotometric method for quantification of C-phycocyanin in the cyanobacterium Aphanizomenon flos-aquae. Journal of Applied Phycology 35(4):1715–1726. 10.1007/s10811-023-03011-1

Blaen PJ, Brekenfeld N, Comer-Warner S, et al (2017) Multitracer Field Fluorometry: Accounting for Temperature and Turbidity Variability During Stream Tracer Tests. Water Resources Research 53(11):9118–9126. 10.1002/2017WR020815

Carrick HJ, Worth D, Marshall ML (1994) The influence of water circulation on chlorophyll-turbidity relationships in Lake Okeechobee as determined by remote sensing. Journal of Plankton Research 16(9):1117–1135. 10.1093/plankt/16.9.1117

Chang G, Dickey TD (2008) Interdisciplinary sampling strategies for detection and characterization of harmful algal blooms. In: Babin M, Roesler CS, Cullen JJ (eds) Real-time Coastal Observing Systems for Marine Ecosystem Dynamics and Harmful Algal Blooms: Theory, Instrumentation and Modelling. chap 2, p 43–84

Cullen JJ (2008) Observation and prediction of harmful algal blooms. In: Babin M, Roesler CS, Cullen JJ (eds) Real-time Coastal Observing Systems for Marine Ecosystem Dynamics and Harmful Algal Blooms: Theory, Instrumentation and Modelling. chap 1, p 1–41

Dunbabin M, Grinham A, Udy J (2009) An Autonomous Surface Vehicle for water quality monitoring. In: Proceedings of the 2009 Australasian Conference on Robotics and Automation, ACRA 2009

Earp A, Dekker AG, Hanson CE, et al (2011) Review of fluorescent standards for calibration of in situ fluorometers: Recommendations applied in coastal and ocean observing programs. Optics Express 19(27):26,768–26,782. 10.1364/OE.19.026768

Foster GM, Graham JL, Stiles TC, et al (2017) Spatial variability of harmful algal blooms in Milford Lake, Kansas, July and August 2015. Tech. rep., 10.3133/sir20165168.

Foy R (1993) The phycocyanin to chlorophyll a ratio and other cell components as indicators of nutrient limitation in two planktonic cyanobacteria subjected to low-light exposures. Journal of Plankton Research 15(11):1263–1276. 10.1093/plankt/15.11.1263

Haggard BE, Grantz E, Austin BJ, et al (2023) Chlorophyll and phycocyanin raw fluorescence may inform recreational lake managers on cyanobacterial habs and toxins: Lake fayetteville case study. Journal of Contemporary Water Research & Education 177(1):63–71. 10.1111/j.1936-704X.2022.3381.x

Hitz G, Pomerleau F, Garneau ME’, et al (2012) Autonomous inland water monitoring: Design and application of a surface vessel. IEEE Robotics and Automation Magazine 19(1):62–72. 10.1109/MRA.2011.2181771

Ji ZG, Jin KR (2006) Gyres and Seiches in a Large and Shallow Lake. Journal of Great Lakes Research 32(4):764–775. 10.3394/0380-1330(2006)32[764:GASIAL]2.0.CO;2

Jin KR, Ji ZG (2004) Case Study: Modeling of Sediment Transport and Wind-Wave Impact in Lake Okeechobee. Journal of Hydraulic Engineering 130(11):1055–1067. 10.1061/(asce)0733-9429(2004)130:11(1055)

Jin KR, Sun D (2007) Sediment Resuspension and Hydrodynamics in Lake Okeechobee during the Late Summer. Journal of Engineering Mechanics 133(8):899–910. 10.1061/(ASCE)0733-9399(2007)133:8(899)

Jin KR, Ji ZG, Hamrick JH (2002) Modeling Winter Circulation in Lake Okeechobee, Florida. Journal of Waterway, Port, Coastal, and Ocean Engineering 128(3):114–125. 10.1061/(asce)0733-950x(2002)128:3(114)

Matson PG, Boyer GL, Bridgeman TB, et al (2020) Physical drivers facilitating a toxigenic cyanobacterial bloom in a major Great Lakes tributary. Limnology and Oceanography 65(12):2866–2882. 10.1002/LNO.11558

Phlips EJ, Badylak S, Nelson NG, et al (2020) Hurricanes, El Niño and harmful algal blooms in two sub-tropical Florida estuaries: Direct and indirect impacts. Scientific Reports 10(1):1–12. 10.1038/s41598-020-58771-4

Rousso BZ, Bertone E, Stewart R, et al (2022) Chlorophyll and phycocyanin in-situ fluorescence in mixed cyanobacterial species assemblages: Effects of morphology, cell size and growth phase. Water Research 212:118,127. 10.1016/J.WATRES.2022.118127

Ruddick K, Lacroix G, Park Y, et al (2008a) Ecosystem dynamics, harmful algal blooms and operational oceanography. In: Babin M, Roesler CS, Cullen JJ (eds) Real-time Coastal Observing Systems for Marine Ecosystem Dynamics and Harmful Algal Blooms: Theory, Instrumentation and Modelling. chap 14, p 527–560

Ruddick K, Lacroix G, Park Y, et al (2008b) Overview of Ocean Colour: theoretical background, sensors and applicability for the detection and monitoring of harmful algae blooms (capabilities and limitations). In: Babin M, Roesler CS, Cullen JJ (eds) Real-time Coastal Observing Systems for Marine Ecosystem Dynamics and Harmful Algal Blooms: Theory, Instrumentation and Modelling. chap 9, p 330–383

Stauffer BA, Bowers HA, Buckley E, et al (2019) Considerations in harmful algal bloom research and monitoring: Perspectives from a consensus-building workshop and technology testing. Frontiers in Marine Science 6(JUL):1–18. 10.3389/fmars.2019.00399

Stockwell JD, Doubek JP, Adrian R, et al (2020) Storm impacts on phytoplankton community dynamics in lakes. Global Change Biology 26(5):2756–2784. 10.1111/gcb.15033

Urquhart EA, Schaeffer BA, Stumpf RP, et al (2017) A method for examining temporal changes in cyanobacterial harmful algal bloom spatial extent using satellite remote sensing. Harmful Algae 67:144–152. 10.1016/J.HAL.2017.06.001

Wachnicka A, Jones P, Marchio D, et al (2022) Appendix 8B-3 : Results from Water Year 2021 Expanded Lake Okeechobee Phytoplankton and Water Quality Monitoring Program. Tech. rep., South Florida Water Management District, West Palm Beach, FL

Welch Z, Zhang J, Jones P, et al (2019) Chapter 8B : Lake Okeechobee Watershed Annual Report. Tech. rep.

Wells ML, Trainer VL, Smayda TJ, et al (2015) Harmful algal blooms and climate change: Learning from the past and present to forecast the future. Harmful Algae 49:68–93. 10.1016/J.HAL.2015.07.009

